# piRPheno: a manually curated database to prioritize and analyze human disease related piRNAs

**DOI:** 10.1101/2020.10.09.334219

**Authors:** Wenliang Zhang, Song Wu, Haiyue Zhang, Wen Guan, Binghui Zeng, Yanjie Wei, Godfrey Chi-Fung Chan, Weizhong Li

## Abstract

Many studies have uncovered that piRNAs (PIWI-interacting RNA) are associated with a broad range of diseases and might be a novel type of biomarkers and targets for precision medicine. However, public resource of high-quality curated human disease-associated piRNAs remains unavailable. Therefore, we developed the piRPheno (http://www.biomedical-web.com/pirpheno) database to provide an up-to-date, interactive and extensible data reference for human piRNA-disease associations. piRPheno includes 9057 experimentally supported associations between 474 piRNAs and 204 diseases through a manual curation of publications. To prioritize the piRNA-disease associations, each association in piRPheno is assigned with a confidence score and clinical correlations based on the experimentally supported evidences. piRPheno is freely available with user-friendly interface and novel applications to enable easy exploration and analysis of the human disease related piRNAs.

## INTRODUCTION

PIWI-interacting RNAs (piRNAs) are an animal-specific class of small silencing RNAs with 21-35 nucleotides in length (Ozata et al., 2019), distinct from microRNAs (miRNAs) and small interfering RNAs (siRNAs). piRNAs bear 2’-O-methyl-modified 3’ termini and guide PIWI-clade Argonautes (PIWI proteins), while miRNAs and siRNAs bear AGO-clade proteins involving in gene silencing pathway (Ozata et al., 2019). piRNAs can guide PIWI proteins to cleave target RNAs, methylate DNA (Kuramochi-Miyagawa et al., 2008), and promote heterochromatin assembly. Moreover, due to the architecture of piRNA signaling pathways, piRNAs are able to regulate expression of conserved host genes, and also provide adaptive immunity and sequence-based immunity (Ernst et al., 2017; Ozata et al., 2019).

With the advances of high throughput sequencing technologies and bioinformatics methods, many piRNAs have been identified and thus the piRNAs data are accumulating rapidly into computational resources, such as piRBase (Wang et al., 2019; Zhang et al., 2014), piRNABank (Sai and Agrawal, 2008), piRNA cluster (Rosenkranz, 2016), piRNAQuest (Sarkar et al., 2014), COMPSRA(Li et al., 2020), piRTarBase (Wu et al., 2019), PingPongPro (Uhrig and Klein, 2019), pirScan (Wu et al., 2018), and IsopiRBank (Zhang et al., 2018). Most of these resources focus on systematically integrating various piRNA associated data to support piRNA functional analysis, biological annotations, and expression profiling. Recently, as piRNAs are implicated in transposon and host gene regulation, many studies have uncovered that piRNA dysfunctions are associated with a broad range of human diseases, such as various cancers (Lee et al., 2016; Mei et al., 2015; Moyano and Stefani, 2015), nervous system disorders (Millan, 2017; Qiu et al., 2017; Roy et al., 2017), and reproductive system disease (Hong et al., 2016).

piRNAs could be a novel type of potential biomarkers and targets for human disease diagnosis, therapy, and prognosis (Millan, 2017; Moyano and Stefani, 2015; Romano et al., 2017). Public resource of manually curated human disease-associated piRNAs remains unavailable. Therefore, in the end of 2018, we started to develop the piRPheno database, which manually curated piRNA-disease phenotype association data from publications. Currently, piRPheno provides 9057 experimentally supported associations between 474 piRNAs and 204 human diseases. To prioritize the piRNA-disease associations, each association in piRPheno is assigned with a confidence score and clinical correlation base on the experimentally supported evidences. In order to enable users exploration and application of the piRNA-disease association data easily, piRPheno (http://www.biomedical-web.com/pirpheno/) provides user-friendly interface and novel visualizations to prioritize and analyze disease related piRNAs online.

## RESULTS

### Data contents

piRPheno provides 9057 experimentally supported associations among 474 piRNAs, 204 disease phenotypes, 26 targets, 16 pathways and 2 treatments. 605 of the associations were manually curated from more than 200publications and 8470 of them were derived by using the disease parent-child relationships in the Experimental Factor Ontology (EFO) resource (Malone et al., 2010). The methods of data curation and new association derivation are detailed in the “Materials and methods” section (Fig. 1). For piRNAs annotation, 97.5 % (462/474) of piRNAs are consistently and comprehensively annotated by piRBase (Fig. 1). For diseases annotation, all of diseases in piRPheno are annotated by the EFO resource (Fig. 1). The piRPheno database covers 6 disease subtypes associated with piRNAs dysregulation, including neoplasm, nervous system disease, reproductive system disease, respiratory system disease, skeletal system disease, and immune system disease. Other than piRNA dysregulation in expression dysregulation, piRPheno also offers 7 single nucleotide polymorphisms (SNPs) on piRNAs are associated with the risk of cancers.

**FIGURE 1.**
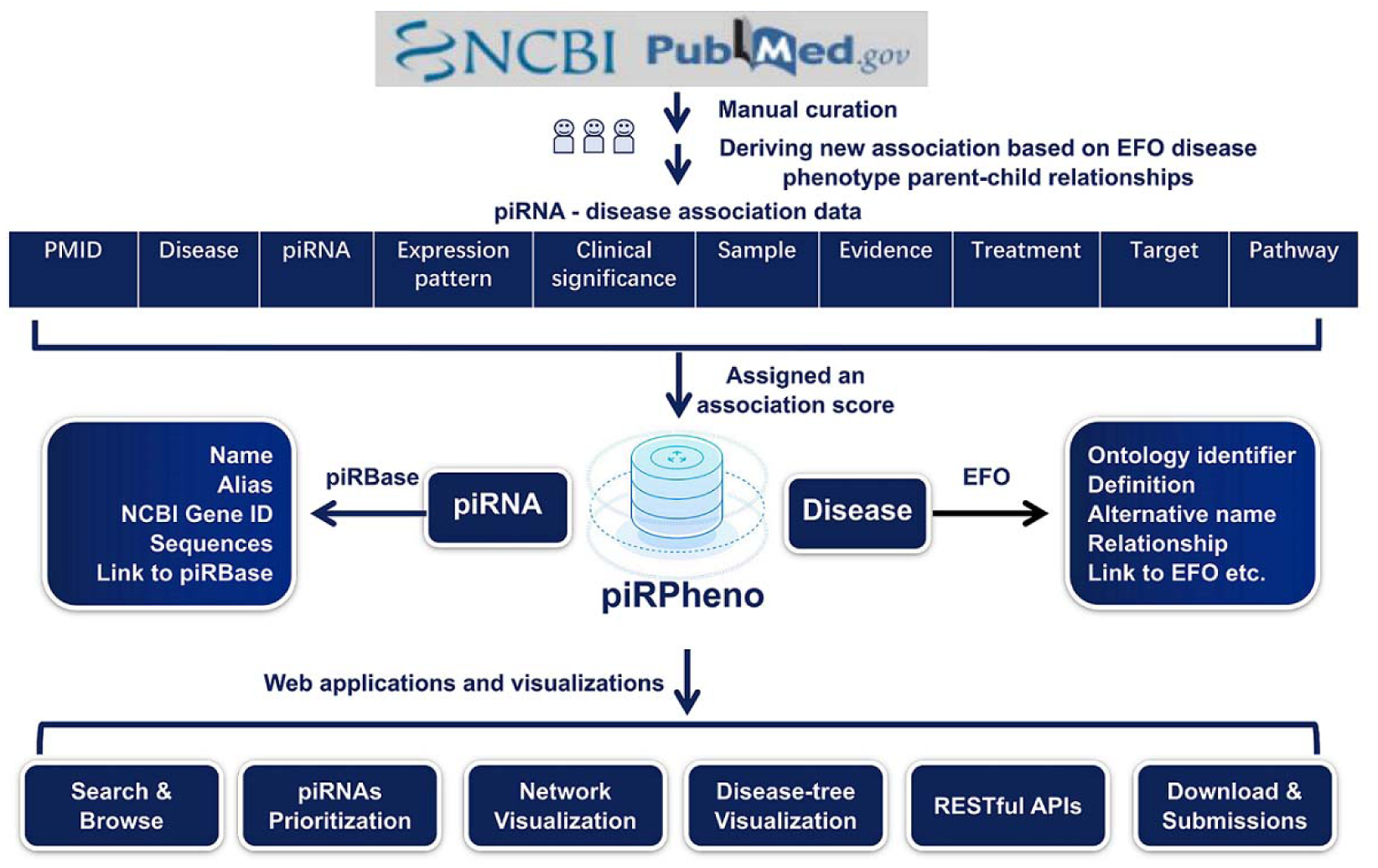
The data curation and annotation framework of piRPheno.

### Search and Browse

The piRPheno database provides user friendly, open access web interfaces and applications to enable users to search, browse, analyze, and prioritize the piRNA-disease association data, as well as to download and submit new associations for further integration (Fig. 1).

To promptly prioritize the piRNA-disease phenotype associations, the search and browse applications were developed in piRPheno. The applications allow users to quickly prioritize piRNA-disease associations through retrieving piRNA and disease phenotype. The search application facilitates smart assistance with keyword tips of expected piRNA and disease phenotype. The prioritizing association data is shown in a brief table, showing key information of association identifiers (IDs), piRNAs, disease phenotypes with ontology identifiers in EFO, confidence scores, and correlations (Fig. 2A). In addition, the prioritizing data allows sorting by confidence scores and filtering by specific piRNA and disease phenotype (Fig. 2A). Moreover, the prioritizing data of a disease search can be optionally visualized in word-cloud diagrams (Fig. 2B), while the prioritizing data of a piRNA search can be optionally visualized in disease-tree and disease-network diagrams (Supplemental_Fig_S1.pptx). Larger sizes and more central locations of the symbols in the word-cloud diagrams indicate higher confidence scores between the piRNAs and disease phenotypes (Fig. 2B). Furthermore, the association IDs in the table, the piRNAs in the word-cloud diagrams, and the circle nodes in the disease-tree diagrams link to further information of the association, piRNA, disease phenotype, and the supporting evidences in publications (Supplemental_Fig_S2.pptx). External links to other reference resources, such as piRBase, EFO, and the NCBI PubMed, and Gene database are also provided (Supplemental_Fig_S2.pptx).

**FIGURE 2.**
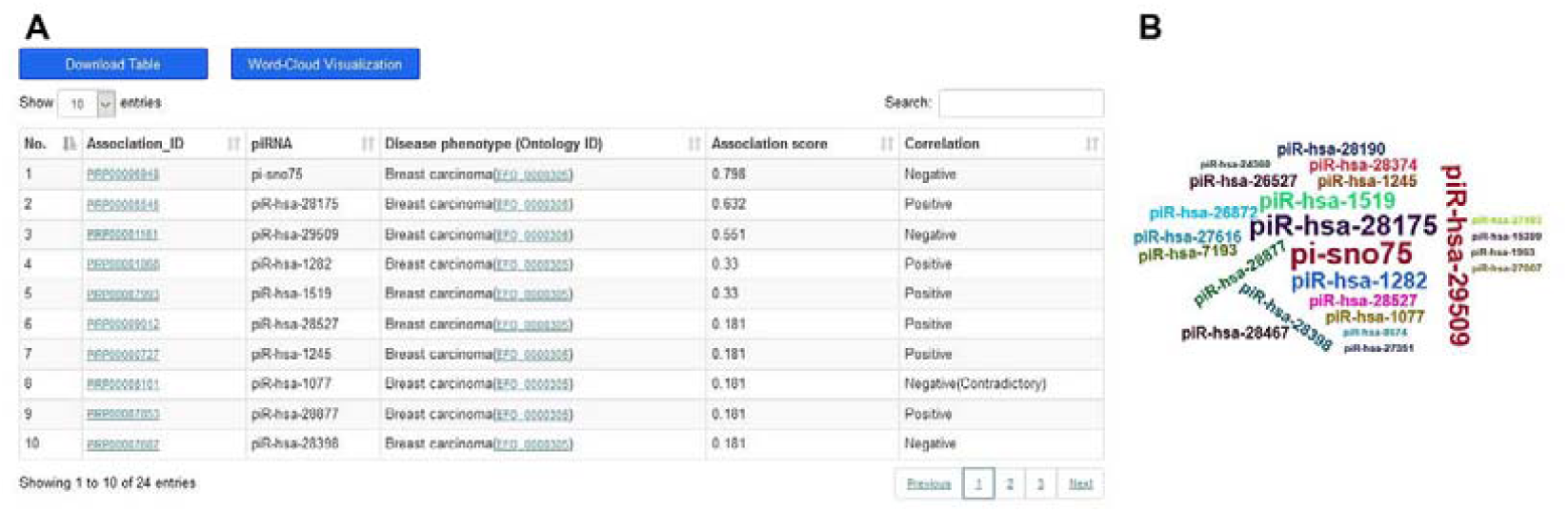
The search and word-cloud application interfaces in piRPheno. (A) Searching results shows breast carcinoma related information in a tabular profile and provides a tabular profile to prioritize piRNAs dysregulation. The “Positive” means that the piRNAs are positively associated with the disease phenotypes, while the “Negative” means that the piRNAs are negatively associated with the disease phenotypes. The “Contradictory” means the piRNA-disease associations with conflicting evidences from different publications. (B) A word-cloud diagram displays the prioritized breast carcinoma related piRNAs. Larger sizes and more central locations of the piRNAs indicate a higher confidence score between the piRNAs and breast carcinoma.

### piRNAs prioritization on disease related piRNAs datasets

A typical case-control microarray assay and piRNA sequencing can find hundreds of significant piRNA dysregulations, but identifying their clinical significance remains challenging. For example, Chu et al investigated and shown that the expression levels of 106 piRNAs were significantly up-regulated and 91 were significantly down-regulated in bladder cancer tissues compared with their corresponding adjacent tissues (Chu et al., 2015). However, how to promptly identify and prioritize the experimentally validated bladder cancer-related piRNAs from these large-scale piRNAs is not a trivial task. To copy with this challenge, a piRNAs prioritization application was developed in piRPheno to analyze and prioritize experimental validated disease phenotype related piRNAs from a set of piRNAs (Fig. 3). We upload 197 piRNAs with bladder cancer phenotype in the piRNAs prioritization application. The application completed the analysis in a few seconds and shown that the piRNA most significantly associated with bladder cancer is piR-hsa-24274. The result table also allows data sorting based on confidence scores and data filtering by specific piRNA (Fig. 3), and it provides links to further webpages for detailed information (Supplemental_Fig_S2.pptx).

**FIGURE 3.**
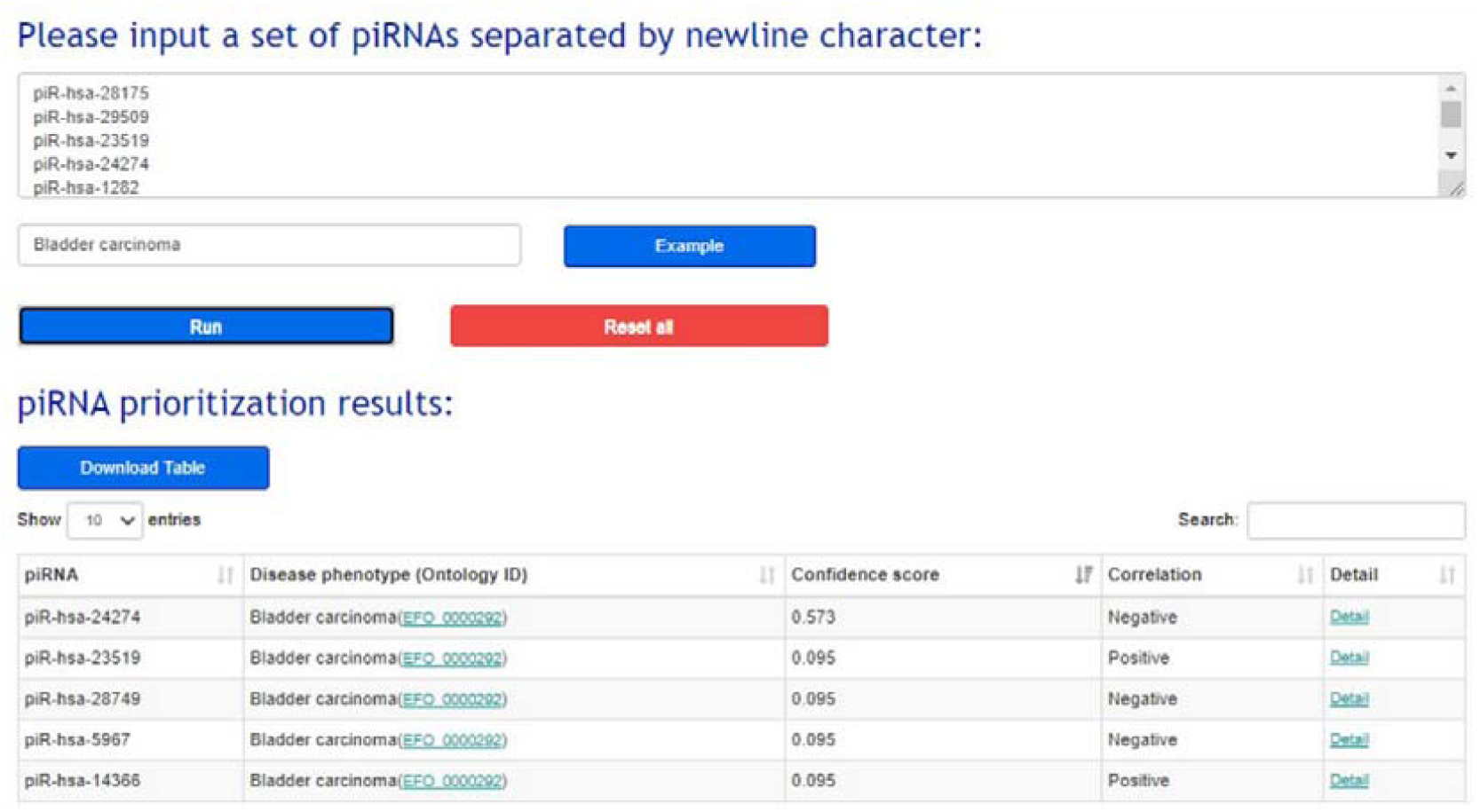
piRNAs prioritization application to promptly prioritize disease related piRNAs from large scale dataset.

### Network visualization to explore the relationships between piRNAs and disease phenotypes

A network visualization application was developed in piRPheno to explore the relationships between piRNAs and disease phenotypes. The application allows user to input a set of piRNAs and disease phenotypes, and to generate interaction networks to display the association data. For example, we entered “breast cancer, lung cancer” in the input box and generated an interaction network to explore the relationships between the two cancers (Fig. 4A). The network clearly indicates that piR-hsa-1077 (Fig. 4A, blue box) is positively associated with the risk of lung cancer, but it is negatively associated with breast cancer and with conflicting evidence from different publications. Similarly, we entered “piR-hsa-8674, piR-hsa-1963” in the input box and generated an interaction network to explore the relationships between the two piRNAs (Fig. 4B). Interestingly, the network clearly indicates that the two piRNAs are both associated with cancer, but piR-hsa-8674 is positively associated with lung cancer and piR-hsa-1963 is negatively associated with colorectal cancer (Fig. 4B).

**FIGURE 4.**
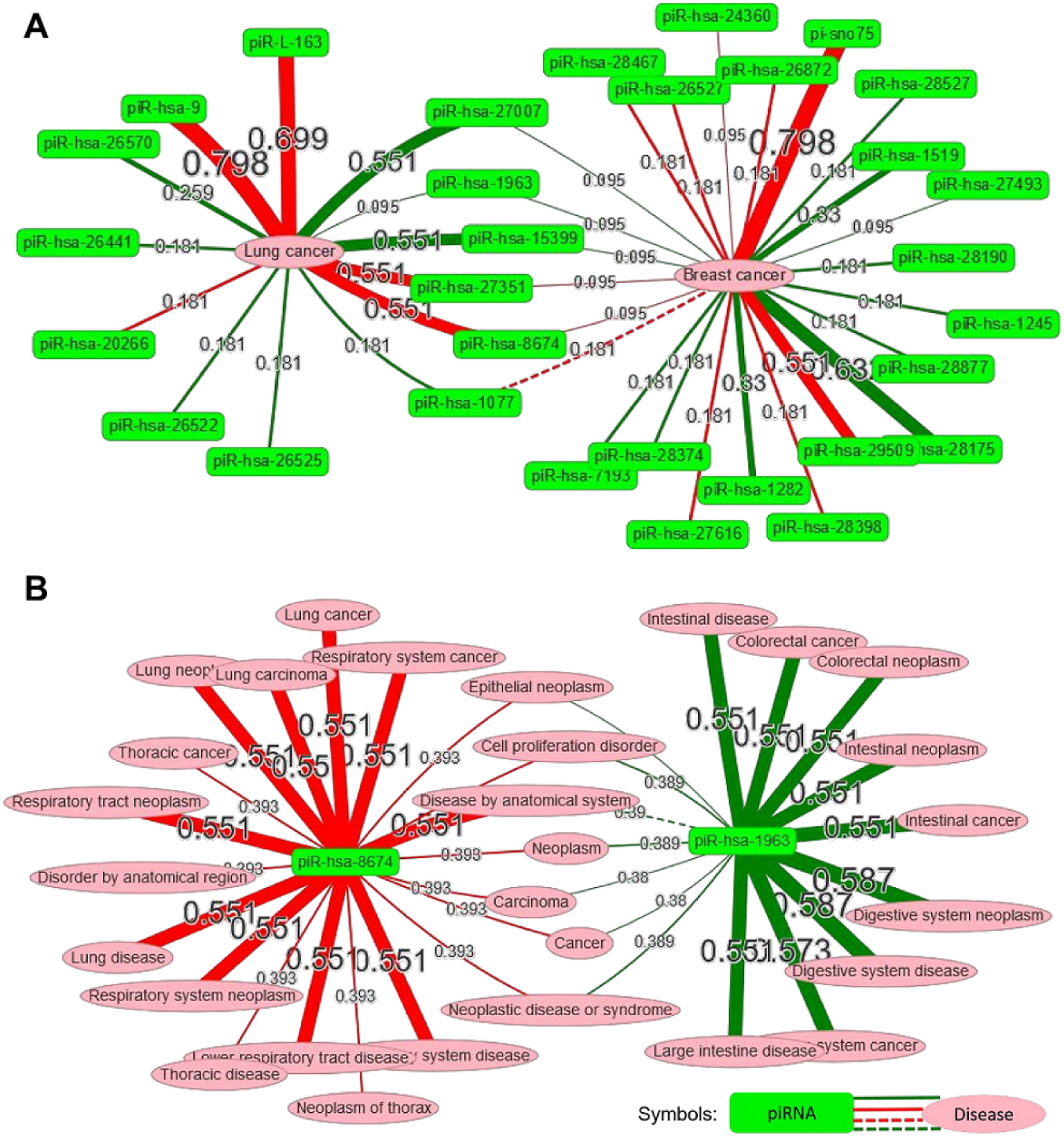
Network visualization to explore the relationships between piRNAs and disease phenotypes. (A) An interaction network to explore the relationships among breast cancer, lung cancer and their related piRNAs. (B) An interaction network to explore the relationships among piR-hsa-8674, piR-hsa-1963 and their related diseases. The red lines indicate that the piRNAs are positively associated with the disease phenotypes, while the green lines indicate that the piRNAs are negatively associated with the disease phenotypes. The dash lines indicate the piRNA-disease associations with conflicting evidences from different publications.

### Data access and submission

piRPheno provides web service APIs for programmatically access of the association data. The resulted data through the APIs are available in the universal JSON formats. Documentation for the use of APIs is available on the “web service” webpage. In addition, all association data in piRPheno is freely available to downloaded and used. Moreover, piRPheno encourages users to submit their new piRNA-disease association data for future data integration. The submitted records will be checked by our professional curators and approved by our submission review committee for the future release. Furthermore, a detailed tutorial is available on the ‘Help’ webpage.

## DISCUSSION

As many studies uncovered that piRNA dysfunctions are associated with a broad range of human diseases, piRNAs is becoming a novel type of potential biomarkers and targets for human disease diagnosis, therapy, and prognosis. Recently, many piRNAs have been identified and several computational resources (Li et al., 2020; Uhrig and Klein, 2019; Wang et al., 2019; Wu et al., 2019; Wu et al., 2018) have been developed to systematically integrate various piRNA associated data to support piRNA functional analysis. Compared with these resources, our piRPheno database not only aims to provide comprehensive and up-to-date data of piRNAs-disease phenotypes association, but also provides novel web applications to analyze and prioritize disease related piRNAs.

The latest update of piRBase (Wang et al., 2019) has collated piRNA-cancer associations from cancer related publications. Compared with piRBase, our piRPheno database not only manually curates piRNA-disease associations from publications, but also derives new associations from the manually curated associations by using the EFO parent-child relationship data. The number of associations in piRPheno is approximately 40-fold of those in piRBase (9057vs. 227). In addition, compared with the latest piRBase, each association in piRPheno is assigned with a confidence score and a clinical correlation base on the experimentally supporting evidences to prioritize and interpret the RNA dysregulation. Importantly, piRPheno provides several novel applications and visualizations to enable easy identification of piRNA dysregulation associated with disease phenotypes for disease diagnosis and therapeutic development, including piRNAs prioritization, disease-tree, word-cloud visualization, and network visualizations.

The piRPheno is updated every 6 months to include new association data and applications. We plan to enrich new association data by analyzing multi-omic data in TCGA (Weinstein et al., 2013) and ICGC (Hudson et al., 2010), and integrate novel bioinformatic tools for further analyzing the piRNA-disease associations in piRPheno.

## MATERIALS AND METHODS

### Data collection and annotation

As previously described (Li et al., 2014; Ning et al., 2016; Zhao et al., 2018), to obtain the all available publications describing the associations between piRNAs and human diseases, we made a query in the National Center for Biotechnology Information (NCBI) PubMed database with the keywords of “(((Piwi-interacting RNA[Title/Abstract] OR Piwi interacting RNA[Title/Abstract] OR piRNA[Title/Abstract]))) NOT review[Publication Type] AND (Humans[Mesh])”. The query resulted in more than 200 publications (before November 2018). We downloaded all of these publications and extracted experimentally supported piRNA-disease association data by manually curation from these publications. Researchers were assigned to double-check all of the collected piRNA-disease associations. In this step, we extracted the piRNA symbol, disease name, experimental evidence, samples, NCBI PubMed ID (PMID), dysfunction status, direct targets, pathway, and treatment (Fig. 1). The clinical significances of piRNA dysfunctions and the experimentally supported evidence levels are also assigned for the piRNA-disease associations (Fig. 1). The clinical significances of piRNA dysregulations are consistently assigned to four status including decreasing risk, increasing risk, decreasing risk with good prognosis and increasing risk with poor prognosis. The assignment of experimentally supported evidence levels of each publication for the piRNA-disease association are shown in detail in Table 1.

**Table 1.**
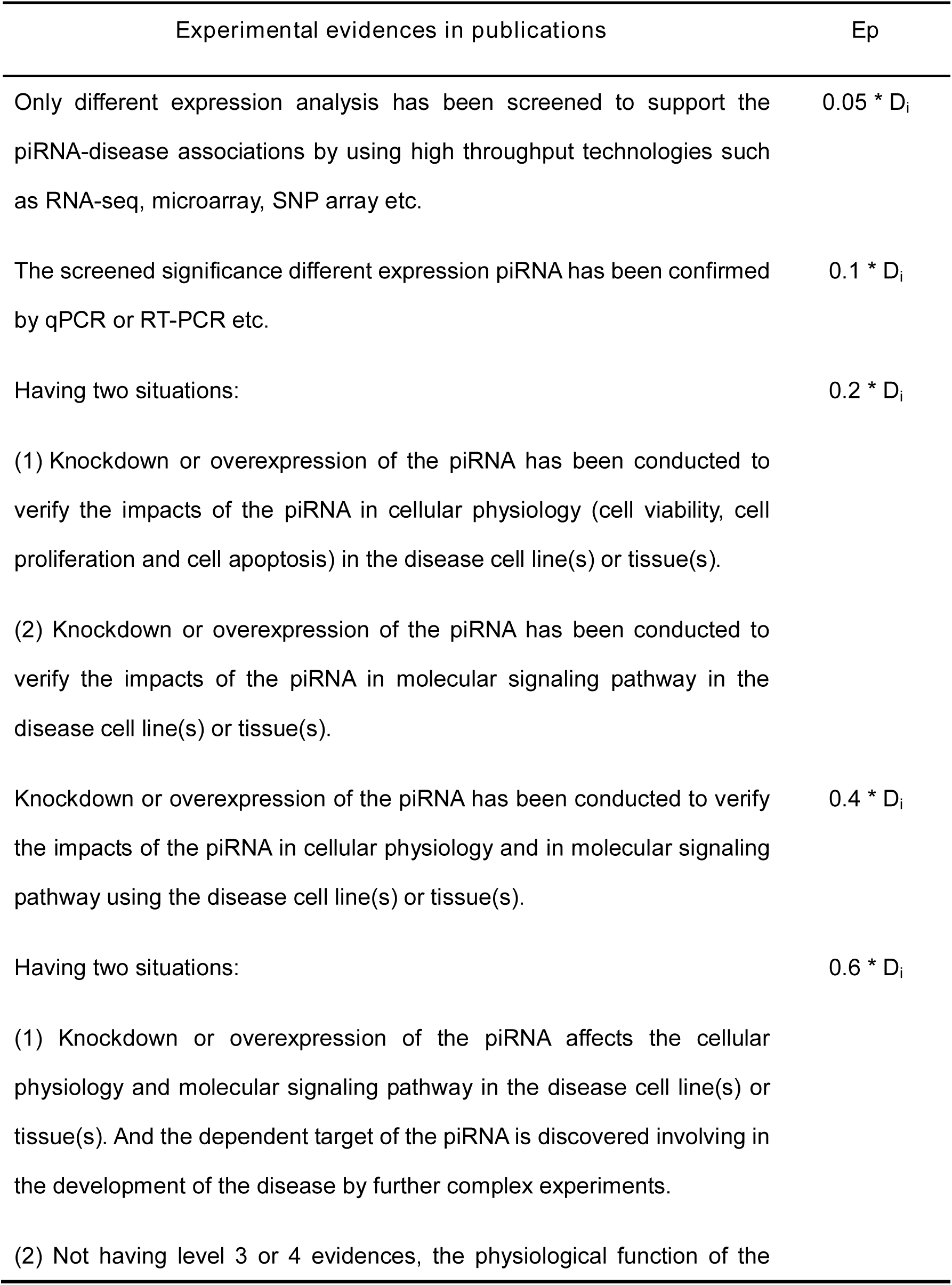
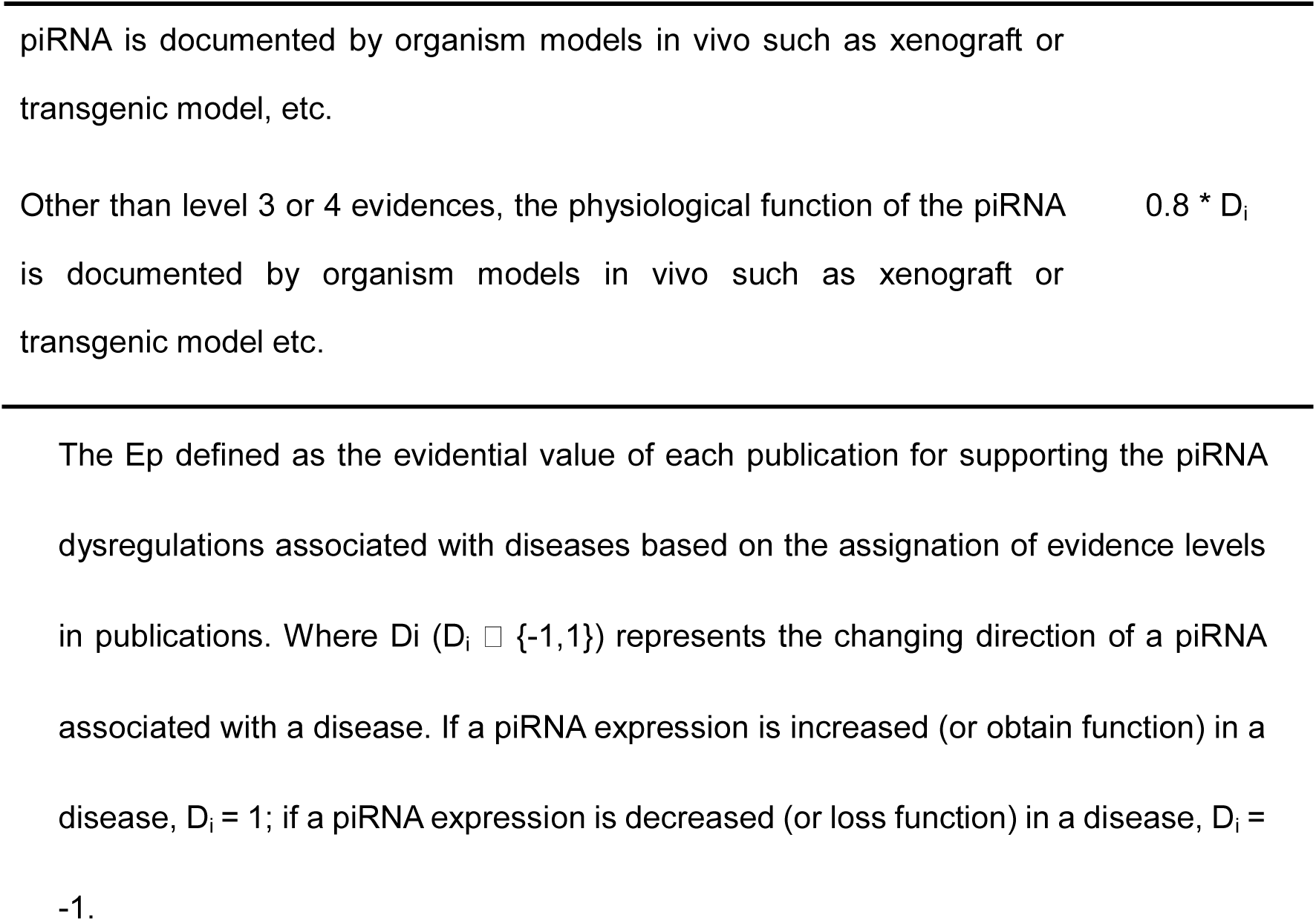
The assignment of experimentally supported evidence levels and the calculation of evidential values

To make the piRNA symbols and disease names consistent with other public databases, the piRBase database offers identifiers, piRNA sequences, alias and links for piRNAs (Fig. 1). Finally, we used a standardized classification scheme, the EFO resource (Malone et al., 2010) to annotate each disease. The annotations of diseases include official disease name, definition, EFO identifier, diseases parent-child relationships, and alterative names (Fig. 1).

### Deriving new piRNA-disease associations

Referred to the Open Target Platform (Koscielny et al., 2017), we also used the EFO parent-child relationship data to derived new piRNA-disease associations, which may not have direct supporting publications, from known piRNA-disease associations with supporting publications (Fig. 1). For example, the non-small cell lung carcinoma and lung adenocarcinoma are both a lung carcinoma. The direct evidence of piRNAs associated to non-small cell lung carcinoma and lung adenocarcinoma are propagated to the higher level of lung carcinomas to allow users to find common piRNAs across groups of related diseases. Other piRNA-disease associations can also be derived based on EFO inferred-by-property classification: disease location (e.g. brain, lung and colon) and disease phenotypes (e.g. azoospermia in male infertility). These two approaches enable driving and propagating new piRNA-disease associations.

### Confidence score

To prioritize and interpret the piRNA dysregulations associated with different diseases in piRPheno, a confidence score for each association is assigned in piRPheno based on two evidential metrics. These evidential metrics include the evidential value in publication (Ep) and the number of publications. The assignment of confidence score consists of three steps:

**Step 1:** In principle, validation experiments of mechanism and functional analyses provide more reliable evidence than throughput expression analyses. Based on the validation experiments in publications, we defined and assigned the experimentally supported evidence levels into six levels, as detailed in Table 1. The evidential value in publication (E_p_) for supporting piRNA-disease association is empirically defined and calculated, as indicated in Table 1. Di (Di∈{-1,1}) in Table 1 represents the changing direction of a piRNA associated with a disease. If a piRNA is increased (or obtain function) in a disease, D_i_ equates to 1; if a piRNA is decreased (or loss function) in a disease, D_i_ equates to −1.

**Step 2:** A large number of publications can enhance the evidential values (Score) for supporting the same piRNA-disease association. To dampen the effect of large number of publications, a harmonic sum function (Hagen, 2008; Koscielny et al., 2017) was used to account Score and abs_Score. The Score and abs_Score are respectively calculating as following equation:

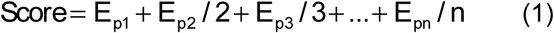

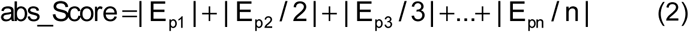

As indicated in equation (1) and (2), “*n*” is the total number of supporting publications, and *E*_*p1*_, *E*_*p2*_, *E*_*p3*_, …, *E*_*pn*_ are the sorted evidential values of different supporting publications in descending order. The Score of an association less than zero indicates that the piRNA dysfunction is negatively associated with the development of disease, and thus the clinical correlation of the association was assigned as “Negative”. On the contrary, the Score of an association greater than zero indicates that the piRNA dysfunction is positively associated with the development of disease, and thus the clinical correlation of the association was assigned with “Positive”. In addition, if the absolute of Score (|Score|) of an association is less than the abs_Score of the association, the clinical correlation of the association was assigned with “Contradictory”. The assignation of “Contradictory” means that the association have conflicting evidence supported.

**Step 3:** The Score above was normalized to limit the range of confidence score from 0 to 1.0.

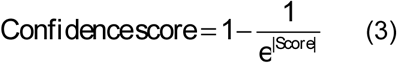

In equation [3], ‘e’ represents the natural constant e.

### Web implementation

The piRPheno website was built with the technologies of Spring MVC and jQuery AJAX framework. Data in piRPheno were organized into a local MySQL database. The programs for data processing were written in Java. The web interface was built by using JavaScript, HTML5, and CSS3. The D3.js widget (http://d3js.org/d3.v3.min.js) and The vis.js widget (http://www.visjs.org) were implemented to display disease-tree visualization and networks on the webpages, respectively. The web service is deployed to an Apache Tomcat web server..

## DATA DEPOSITION

The data in piRPheno is available at http://www.biomedical-web.com/pirpheno.

## SUPPLEMENTAL MATERIAL

Supplemental material is available for this article at http://www.biomedical-web.com/pirpheno/suppl.jsp.

## ACKNOWLEDGMENTS

This work has been supported by Sanming Project of Medicine (Shenzhen) [SZSM201911016], the National Key R&D Program of China [2016YFC0901604 & 2018YFC0910401], the National Natural Science Foundation of China [31771478], the Fundamental Research Funds for the Central Universities, Sun Yat-sen University (No.19ykpy86), and the China Postdoctoral Science Foundation (No.2020M673023).

